# Prediction of 8-state protein secondary structures by 1D-Inception and BD-LSTM

**DOI:** 10.1101/871921

**Authors:** Aminur Rab Ratul, Marcel Turcotte, M. Hamed Mozaffari, WonSook Lee

## Abstract

Protein secondary structure is crucial to create an information bridge between the primary structure and the tertiary (3D) structure. Precise prediction of 8-state protein secondary structure (PSS) significantly utilized in the structural and functional analysis of proteins in bioinformatics. In this recent period, deep learning techniques have been applied in this research area and raise the Q8 accuracy remarkably. Nevertheless, from a theoretical standpoint, there still lots of room for improvement, specifically in 8-state (Q8) protein secondary structure prediction. In this paper, we presented two deep learning architecture, namely 1D-Inception and BD-LSTM, to improve the performance of 8-classes PSS prediction. The input of these two architectures is a carefully constructed feature matrix from the sequence features and profile features of the proteins. Firstly, 1D-Inception is a Deep convolutional neural network-based approach that was inspired by the InceptionV3 model and containing three inception modules. Secondly, BD-LSTM is a recurrent neural network model which including bidirectional LSTM layers. Our proposed 1D-Inception method achieved 76.65%, 71.18%, 76.86%, and 74.07% Q8 accuracy respectively on benchmark CullPdb6133, CB513, CASP10, and CASP11 datasets. Moreover, BD-LSTM acquired 74.71%, 69.49%, 74.07%, and 72.37% state-8 accuracy after evaluated on CullPdb6133, CB513, CASP10, and CASP11 datasets, respectively. Both these architectures enable the efficient processing of local and global interdependencies between amino acids to make an accurate prediction of each class is very beneficial in the deep neural network. To the best of our knowledge, experiment results of the 1D-Inception model demonstrate that it outperformed all the state-of-art methods on the benchmark CullPdb6133, CB513, and CASP10 datasets.

## Introduction

Proteins are the vastly complex substance, which is the basis of any living organism and involved in every part of cellular life. Proteins perform a colossal number of functions such as DNA replication, transporting molecules across membranes, giving structures to organisms, catalyzing chemical reactions, run the immune system, and regulating oxygenation. Proteins are long-chain molecules made up of hundreds and thousands of 20 different kinds of smaller units called amino acids. It is necessary to know the protein structure to understand the functionality of proteins. Sometimes it is essential to know protein 3D structures to identify the protein functions at a molecular level. Reliably and accurately predicting protein 3D structure form sequences of proteins is one of the most challenging issues in computational biology [1]. Protein secondary structure prediction is a vital step towards to predict protein tertiary (3D) structure [2], prediction of protein disorder [3], and solvent accessibility prediction [4].

In protein secondary structure (PSS), there are two regular PSS types: alpha-helix (H) and beta-strand (E), and one irregular PSS state: coil region (C). In protein secondary structure prediction (PSS) together, predict these three states, or the accuracy of these states is known as Q3 accuracy. According to hydrogen-bonding patterns, Sander developed the Dictionary of Secondary Structure of Proteins (DSSP) algorithm to classify protein secondary structure into 8-states [5]. Furthermore, the DSSP algorithm designated helix into three types (G or 3-10 helix, H or α-helix, and I or π-helix), two kinds for strand (E or β-strand, and B or β=bridge), and coil into three states (T or hydrogen-bonded turn, S or high curvature loop, and L or, irregular). The performance measure of these eight states is the Q8 accuracy (the 8-states protein secondary structure classification accuracy).

Previously many methods extensively studied to predict protein secondary structure, including support vector machine techniques (SVM) [6, 7], artificial neural networks [8–10], hidden Markov model [11, 12], and bidirectional recurrent neural network (BRNN) [13–16], and the probability graph models [17, 18]. Because of the advancement of deep learning, recently, most of the protein secondary structure prediction models based on deep learning and provide us with promising outcomes [19, 20–22, 23, 24]. Most of the methods of protein secondary structure prediction (PSS) have been heavily focused on state-3 (Q3) classification. However, predict the 8-classes protein secondary structure is more challenging and complex, and the 8-categories of PSS reveals more detailed local information. Additionally, only a few deep learning architectures exist which can build to predict 8-states PSS. Therefore, in this paper, we only focus on to predict 8-categories of protein secondary structure based on the sequences of protein.

Here, in order to achieve the Q8-accuracy of protein, we utilized two different deep learning methods. At first, we establish a one dimensional (1D) deep convolutional neural network (DCNN) based model called 1D-Inception, which inspired by a widely used computer vision model InceptionV3. Next, we proposed another architecture that focuses on a popular recurrent neural network (RNN) LSTM namely, Bidirectional LSTM (BD-LSTM). Furthermore, to training, validation, and testing, we use five public datasets: CullPdb 6133, CullPdb 6133 filtered, CB513, Casp10, and Casp11.

The rest of the paper organized as follows: in section 2, discuss several related works. The dataset and methodology demonstrated in section 3. Section 4 illustrated the performance analysis of the experiment. Finally, the conclusion and discussion displayed in section 5.

## Related Works

In 1951, when Pauling and Corey [25, 26] predicting hydrogen-bonding patterns from α-helices and β-sheets that open the door for protein secondary structure research. Although in 1958, the first protein secondary structure (PSS) prediction discovered by X-ray crystallography [27]. The protein secondary structure prediction approach split into three different generations [28]. Secondary structure prediction accomplishes from amino acid residues’s statistical propensities of the protein sequence in the first generation [29–31], and the most popular method of this was Chou–Fasman method [31]. The second-generation techniques employed a sliding window of neighboring residues (LIM method [32], GOR method [33]) and diverse theoretical algorithms like nearest neighbor algorithm [34], graph theory [35], neural network based techniques [36], statistical information [33, 37], and logic-based machine learning approach [38]. The third-generation methods mostly based on evolutionary information obtained from the alignment of multiple homologous sequences [39]. The most notable enhancement in protein secondary structure prediction achieved by Position-specific scoring matrix (PSSM) based on PSIBLAST [40] reflects evolutionary information. Many novel computational algorithms have been proposed during this time. For instance, Bayesian or hidden Markov model [41], conditional random fields for combined prediction [42], and support vector machines [43]. Among all of these latest models, deep learning based approaches provide superior reported accuracy [20, 23, 44, 21, 22]. Specifically, in this period, the Q3 accuracy of protein secondary structure reached above 80%. Although the traditional methods are not able to produce a satisfactory Q8 accuracy rate because classify 8-categories of protein secondary structure is more complicated and challenging [2, 19, 45].

In recent times, deep neural network (DNN) models have become established tools for the representation learning of many distinct data [46–48]. By incorporating evaluation information deep neural networks (DNNs) also attained significant enhance accuracy on the state-8 (Q8) protein secondary structure prediction [19]. Wang et al. [23] integrating shallow neural network and Conditional Random Fields (CRF) and proposed Deep Convolutional Neural Fields (DeepCNF) architecture. The experimental result shows that DeepCNF can achieve almost 84% Q3 accuracy and 72% Q8 accuracy on the CASP, and CAMEO test proteins, respectively. Based on a semi-random subspace method (PSRSM) and data partition in [44], Ma et al. presented a novel architecture that utilized Support Vector Machines (SVMs) as the base classifier. They achieved 86.38%, 84.53%, 85.51%, 85.89%, 85.55%, and 85.09% Q3 accuracy respectively on 25PDB, CB513, CASP10, CASP11, CASP12, and T100 datasets. A new deep learning architecture suggested by Zhou et al. [20], namely CNNH_PSS, where they were using multi-scale CNN with highway to predict the accuracy of PSS. They accomplished 74.0%, and 70.3% Q8 accuracy on CullPdb6133, and CB513, respectively. Guo et al. [21] proposed the DeepACLSTM model to predict 8-classes secondary structure from profile features and sequence features. In DeepACLSTM, they combined ACNNs (asymmetric convolutional neural networks) with BLSTM (bidirectional long short-term memory) neural networks together and found 74.2%, 70.5%, 75.0%, 73.0% Q8-accuracy respectively on CullPdb6133, CB513, CASP10, and CASP11. Fang et al. [19] provide a Deep inception-inside-inception (Deep3I) model to predict the Q3 and Q8 accuracy for PSS, and later they implemented this as a software tool called MUFOLD-SS. MUFOLD-SS obtained 70.63%, 76.47%, 74.51%, 72.1% Q8 accuracy respectively on CB513, CASP10, CASP11, and CASP12. Zhang et al. presented a novel method based on a convolutional neural network (CNN), bidirectional recurrent neural network, and residual network to enhance the prediction accuracy for the 8-state protein secondary protein structure (Q8) [49], and attained 71.4% accuracy on the benchmark CB513 dataset. In [50], Busia et al. suggested an ensemble method consisting of various popular deep neural architectures-DenseNet, ResNet, and Inception with BatchNomalization-to predict the Q8 accuracy of PSS and reaches 70.6% state-8 accuracy on CB513 dataset.

## Datasets and Methodology

### Datasets

Here, we utilize five different datasets, namely, CullPdb 6133, CullPdb 6133 filtered, Cb513, Casp10, and Casp11. Among these five datasets CullPdb 6133, and CullPdb 6133 filtered for training. Furthermore, CB5133, Casp10, Casp11, and 272 protein sequence of CullPdb 6133 for testing.

#### CullPdb 6133

CullPdb 6133 [51] dataset is a non-homologous protein dataset that is provided by PISCES CullPDB with the familiar secondary structure for protein. This dataset contains a total of 6128 protein sequences, in which 5600 ([0:5600]) protein samples are considered as the training set, 272 protein samples [5605:5877] for testing, and 256 proteins samples ([5877,6133]) regarded as the validation set. Moreover, CullPdb 6133 (non-filtered) dataset has 57 features, such as amino acid residues (features [0:22)), N- and C- terminals (features [31,33)), relative and absolute solvent accessibility ([33,35)), and features of sequence profiles (features [35:57)). We used secondary structure notation (features [22:31)) for labeling. This CullPdb dataset is publicly obtainable from [2].

#### CullPdb 6133 filtered

There exists redundancy between the CB513 dataset and CullPdb 6133 dataset. Therefore, in CullPdb 6133 filtered version 5534 protein sequences created by detaching those sequences whose over 25% resemblance between CB513 and CullPdb6133. In this dataset, we take a 5234 protein sequence randomly picked for training and the remaining 300 protein sequences for validation.

#### CB513

CB513 dataset contains 514 protein sequences, and it is widely utilized to compare the performance of protein secondary structure prediction. Here, we use this dataset for evaluating our testing result after training with CullPdb 6133 filtered. This dataset can be accessed from [2, 52].

#### Casp10 and Casp11

Casp10 and Casp11 [20, 24] respectively contain 123 and 105 protein sequences. Since 1994, the Critical Assessment of protein Structure Prediction or CASP used for protein structure prediction all over the world. The bioinformatics community has hugely exploited these datasets. After training our models with CullPdb6133 filtered, we were using the CASP dataset for testing.

To use the features from CullPdb 6133 and CullPdb 6133, we reshaped these datasets from 6133 proteins x 39900 features and 5534 proteins x 39900 features into 6133 proteins x 700 amino acids x 57 features and 5534 proteins x 700 amino acids x 57 features respectively. Here, 700 indicate the peptide chain, and 57 signify the features for each amino acid. We use amino acid residues and sequence profiles (total of 42 features) for training input. Besides, the output is a protein secondary structure labeling.

### Methodology

In this paper, we implement two deep learning architectures to predict protein secondary structure prediction. The first model based on a one-dimensional (1D) deep convolutional neural network, and for the second model, we use the popular recurrent neural network (RNN) based approach, specifically LSTM.

#### Deep Convolutional Neural Network Model (1D-Inception)

Lately in computer vision InceptionV3 [53] architecture becomes popular after producing high accuracy on ImageNet [54] dataset with less computational cost than many existing popular architectures. We hugely inspired by the InceptionV3 model to build this architecture. We also get an idea from the literature [19] when we tried to create this model for protein secondary structure prediction, although our approach is very much different than this literature.

The basic InceptionV3 network for image recognition has three different parts: 1) Factorizing Convolutions; 2) Auxiliary Classifier; and 3) Systematic Grid Size Reduction. Throughout the whole network, different window size has been used for convolution operations such as 1×1, 3×3, 5×5, 7×7. The 1×1 convolution employed to lessen the dimensionality of the feature map. Additionally, extracting the image features max-pooling operation used in this network.

**Figure 1:**
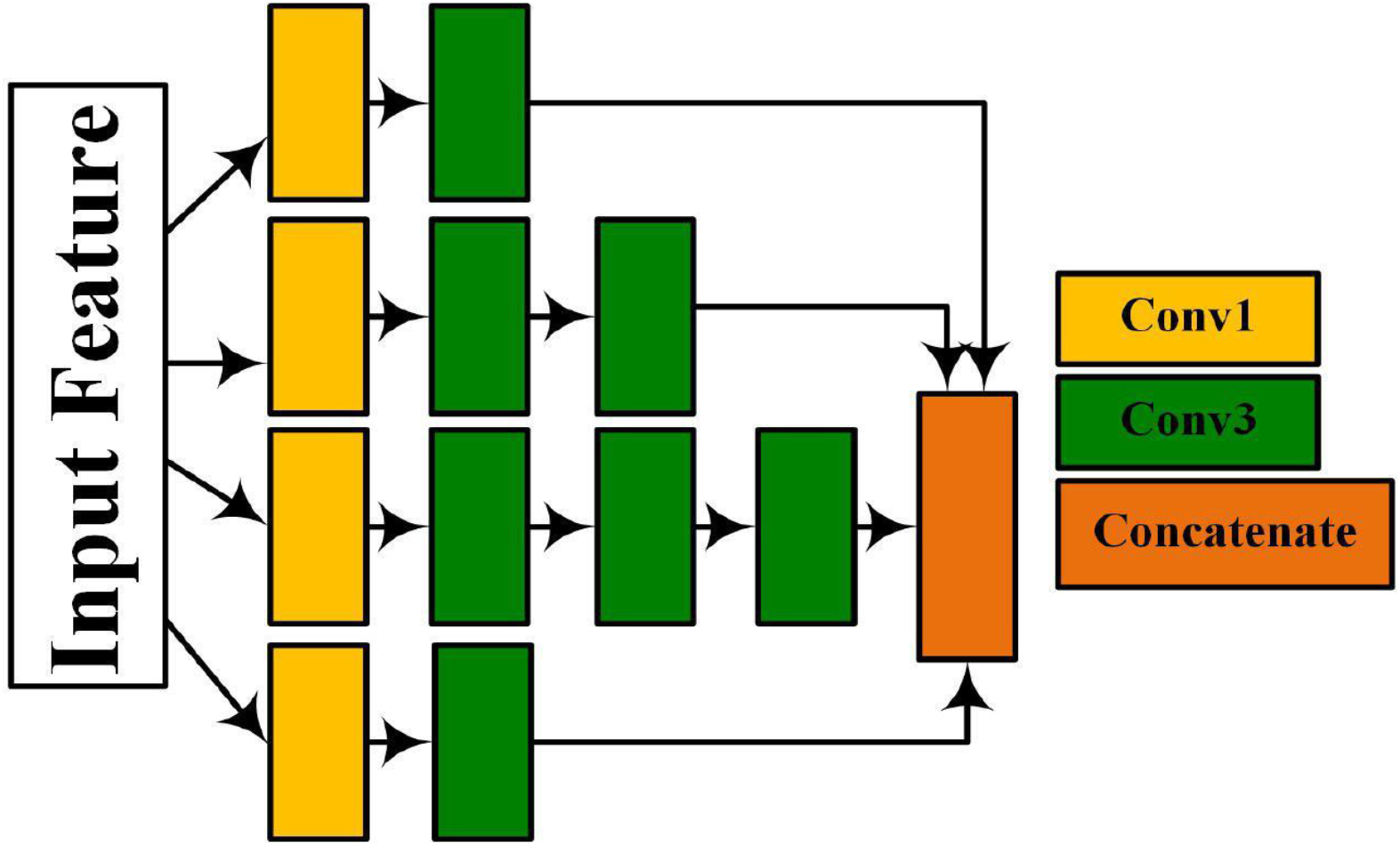
This is a 1D-Inception module. The Yellow rectangle bar means “Conv1”- 1×1 convolution operation, the Green rectangle bar stands for “Conv 3”- 3×3 convolution operation, and the Orange rectangle bar means “Concatenate”- accomplishes feature concatenation.

In this experiment, to predict the secondary structure of the protein, we do not use max pooling because there is a chance to change the sequence length of the protein. There are three separate modules in our networks. In every module, to extract various features from the input sequence, we utilized several parallel convolutional layers in each module. The input of the architecture is *I* × *d* matrix. Here, the length of the input sequence defines as *I*, and *d* is the dimension of vectors. So the value of *I* = 700 and *d* = 42. The output of this network is secondary structure labels. We use four different window sizes throughout the system, like 1×1, 3×3, 5×5, 11×11.

All the convolutional layer in this network is one dimensional, using Rectified Linear Unit (RELU) [55] as the activation function, and *Padding = “same”* which means output has the identical length as input. “Conv 1”, “Conv 3”, “Conv 5”, and “Conv 11” has kernel size 1, 3, 5, and 11 respectively. “Conv3” and “Conv11” has some other operations: 1. To increase the speed of the training process, we use the Batch Normalization [10] that is also working as a regularizer, 2. *dropuout* = 0.25 to prevent the overfitting by randomly dropping 25% neurons from a particular layer.

In the input layer of this architecture, we apply the he_uniform [57] method for random weight initialization. The uniform distribution of this technique lies between –*limit to* + *limit* where 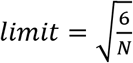 and *N* = *total number of input units in the weight tensor*.

**Figure 2:**
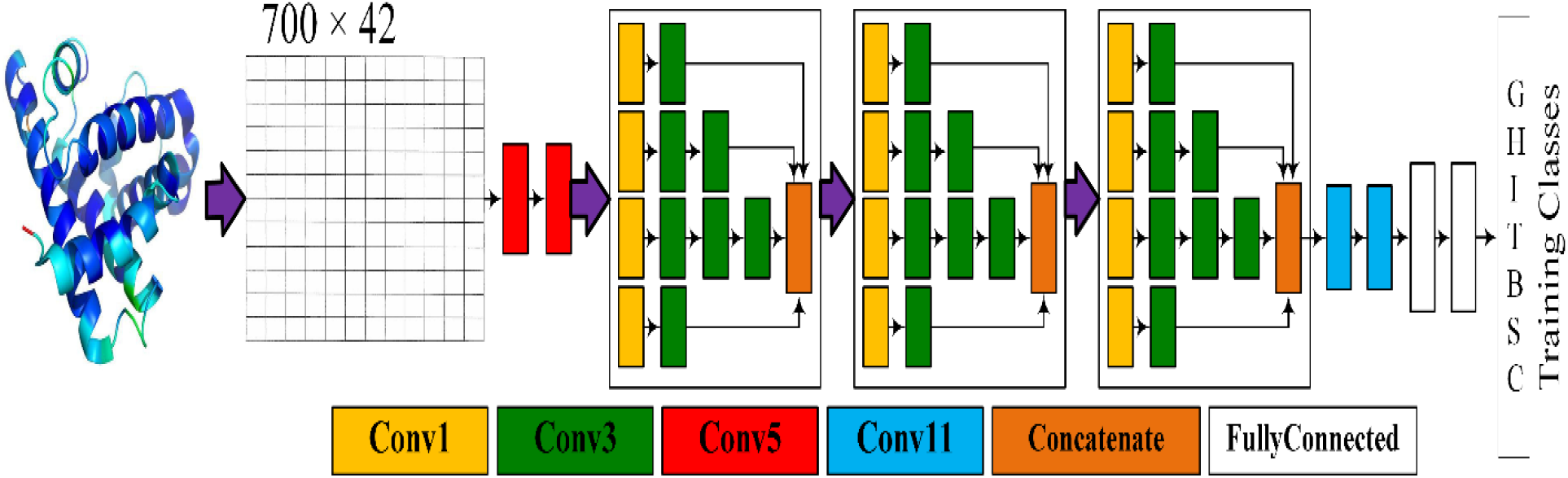
A 1D-Inception network that consists of three 1D-Inception modules. This network has two “Conv 5” blocks followed by three 1D-Inception modules. After that, there are two “Conv 11” blocks with two fully connected layers in the last part of this architecture.

The number of the filters in every convolutional layer in Module A is 256, Module B has 128 number of the filters in its convolutional layer, the number of filters in Module C is 64, “Conv 5” has 512 filters, and “Conv 11” has 128 filters. The last two-layer of this network is fully connected layer has 256 and 8 filters respectively with *dropuout* = 0.25 in between. The first fully connected layer has the RELU activation function, and the last one has the softmax activation function.

#### Bidirectional LSTM Model (BD-LSTM)

To predict protein secondary structure, we further try one recurrent neural network (RNN) based approach. To build this model, we use three Bidirectional LSTM (Long Short Term Memory) layer with seven fully connected layers. To provide the fixed variable length and handle the variable length in the input layer, we use masking with 700 × 21 input matrix length like previously. Then apply three bidirectional LSTM units with 128, 256, and 512 filters, respectively. Furthermore, in these three LSTM layers, we also use “Tanh” as the activation function and *dropuout* = 0.25. Besides, the dropout rate 0.25 throughout the whole network. We concatenate the output of three bidirectional LSTM units and feed the output to the fully connected layers. We add seven fully connected layers one after another, where the first two architecture has 512 filters, the sixth layer has 256 filters, and the last layer has 8 filters for secondary structure labeling. We employ the RELU activation function in all fully connected layers except the last one, where we use the SoftMax activation function. Maintain one to one relationships on input and output in the final layer, we utilized Timedistributed fully connected layer.

**Figure 3:**
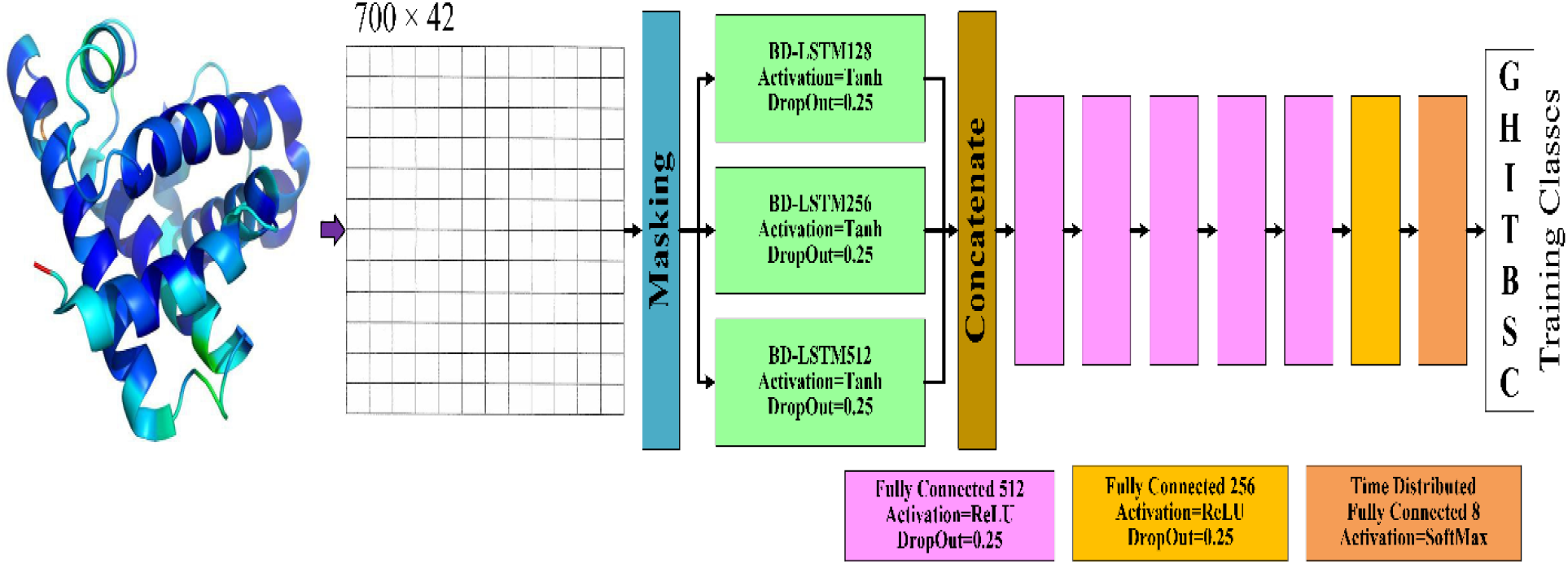
BD-LSTM architecture has three bidirectional-LSTM layers, followed by seven fully connected dense layers.

These two networks were implemented using TensorFlow as backend and Keras as the frontend library. Furthermore, the mini-batch size of these two networks is 32. For the Deep Convolutional model, we take .0002 as our learning rate, and for the LSTM network, we apply .002 as the learning rate. We have utilized Adam optimizer [12] as the optimization function and train both architectures for 125 epochs or iterations. During train our networks, we inspect the validation loss for 10 epochs to see that validation loss is reducing or not. If it does not lessen, then we decrease the learning rate as the factor by (0.1)^5^.

Learning rate would be:

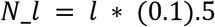

*N_l* = new learning rate; *I* = present learning rate. In addition, the lower bound of the learning rate is 0.5*e* – 5.

Furthermore, we employ “Categorical Crossentropy” as the loss function in both of our models.

## Result

Here, we only focus on the overall accuracy of 8 classes (Q8) of protein secondary structure. In this section, broad experimental outcomes for Q8 protein secondary structure prediction displayed of proposed deep learning several architectures on popular datasets and compared the performance with some state-of-art method. At first, we tried several different combinations for these two proposed architectures to decide what would be the perfect setup. We execute our experiment in two ways and show the evaluation of the proposed architecture.

We use several datasets to evaluate the performance of the architectures. Firstly, train on 6128 protein sequences of CullPdb 6133, validated on 256 protein samples on the same dataset, and tested on the remaining 272 protein sequences. Next, picked 5234 protein sequence randomly for training from CullPdb 6133 filtered dataset, validated on the remaining 300 sequences, and tested on Cb513, Casp10, Casp11. In table 1, we manifest the peak validation outcomes, and test result for several combinations of deep CNN based approaches (InceptionV3 look alike).

**Table 1:** Q8 (State-8) accuracy of 1D-Inception Network with Different Numbers of Modules and best outcomes marked in bold

| 1D Inception: Module Number | Test Accuracy on CullPdb 6133 (%) | Test Accuracy on CB513 (%) | Test Accuracy on CASP10 (%) | Test Accuracy on Casp11 |
| --- | --- | --- | --- | --- |
| 1 | 72.97 | 68.39 | 73.51 | 70.24 |
| 2 | 75.88 | 70.61 | 75.70 | 72.93 |
| 3 | **76.65** | **71.18** | **76.86** | **74.07** |
| 4 | 75.19 | 70.04 | 74.85 | 73.63 |
| 5 | 74.34 | 69.47 | 73.79 | 72.54 |

Moreover, for our second architecture, we also tried several different combinations. We fixed the position of the first three Bidirectional LSTM layers and the last two fully connected layers. However, only add or remove the fully connected layer in between where each layer has 512 filters. In table 2, we displayed the same output for the different combinations for the BD-LSTM based approach.

**Table 2:** Q8 (State-8) accuracy of BD-LSTM Network with Different combination and best outcomes marked in bold

| BD-LSTM Model | Test Accuracy on CullPdb 6133 (%) | Test Accuracy on CB513 (%) | Test Accuracy on CASP10 (%) | Test Accuracy on Casp11 |
| --- | --- | --- | --- | --- |
| Model 1 | 70.62 | 67.87 | 70.33 | 69.50 |
| Model 2 | 72.95 | 68.06 | 72.28 | 70.85 |
| Model 3 | 73.40 | 68.83 | 72.94 | 71.79 |
| Model 4 (Proposed approach) | **74.71** | **69.49** | **74.07** | **72.37** |
| Model 5 | 73.55 | 69.13 | 73.57 | 71.65 |

Combinations of Bidirectional LSTM architecture:

1. Masking+3BD-LSTM+2FC(512 units)+1FC(256 units)+1 TimeDistributed-FC(8 unit)
2. Masking+3BD-LSTM+3FC(512 units)+1FC(256 units)+1 TimeDistributed-FC(8 unit)
3. Masking+3BD-LSTM+4FC(512 units)+1FC(256 units)+1 TimeDistributed-FC(8 unit)
4. **Masking+3BD-LSTM+5FC(512 units)+1FC(256 units)+1 TimeDistributed-FC(8 unit)**
5. Masking+3BD-LSTM+6FC(512 units)+1FC(256 units)+1 TimeDistributed-FC(8 unit)

Protein secondary structure prediction is a vital problem to solve in bioinformatics because of drug discovery [13, 4] and analyzing protein function. There are many state-of-art methods proposed to predict state-8 (Q8) protein secondary structure. In this section, the comparison between our two proposed models and following state of the art architectures.

### SSPro-8

In [60], Pollastri et al. utilized PSI-BLAST-derived profiles and ensemble the Bidirectional RNN model to enhance the state-8 prediction accuracy of protein secondary structure.

### MUFOLD-SS-8

A new deep neural network proposed to map protein secondary structure where Fang et al. [19] introduce a nested inception network. We also use an InceptionV3 look-alike model, but our approach and architecture design is a lot different than this model.

### CNNH_PSS

Zhou et al. [20] proposed a deep learning based approach, namely CNNH_PSS, by employing a multi-scale Convolutional network with the highway technique to predict Q8 PSS.

### DeepACLSTM

In [21], we find a model where Guo et al. provide an asymmetric CNN (ACNN) merge with Bidirectional LSTM to predict protein secondary structure. They use sequence features and profile features to find 8 classes of PSS.

### 2C-BRNN

To improve the accuracy of 8-state PSS Guo et al. [22] proposed hybrid deep learning architectures 2D Convolutional Bidirectional RNN (2C-BRNN) based approach. They used Bidirectional Long Short term Memory and Bidirectional Gated Recurrent Units to construct four different models. Their proposed architectures are: 2DCNN-BGRUs,2DConv-BGRUs,2DCNN-BLSTM, and 2DConv-BLSTM.

### CNF

Wang et al. [61] present a probabilistic model graphic model called conditional neural fields (CNFs) to predict the 8-categories PSS. By this method, they model the complex relationship between sequence features, PSS and utilize the interdependency among PSS kinds of adjacent residues.

### DeepCNF

In [23], Wang et al. extend their CNF model using deep learning techniques to make it more powerful. It is an integration between shallow neural networks and Conditional Random Fields (CRF).

### GSN

To predict protein secondary structure Zhou et al. [2] present a supervised Generative Stochastic Network (GSN) with deep hierarchical representations.

### DCRNN

Li et al. [24] proposed an end-to-end deep neural network that predicts state-8 PSS from integrated local and global contextual features.

### Training and Testing on CullPdb6133 Dataset

First of all, we compare the Q8 accuracy of the protein secondary structure of our proposed models and five other benchmark models, which trained and tested on the CullPdb6133 dataset.

According to table 3, our proposed 1D-Inception model attains the highest Q8 accuracy that exceeds the accuracy of all the benchmark architectures for CullPDB6133. 1D-Inception exhibits 76.65% overall state-8 accuracy, which is .95% more than 2DConv-BLSTM [19] model and 1.45% more than DeepCNF. Furthermore, our other proposed model BD-LSTM provides better accuracy than most of the benchmark models except 2DConv-BLSTM, DeepCNF, and 2DConv-BGRUs.

**Table 3:**
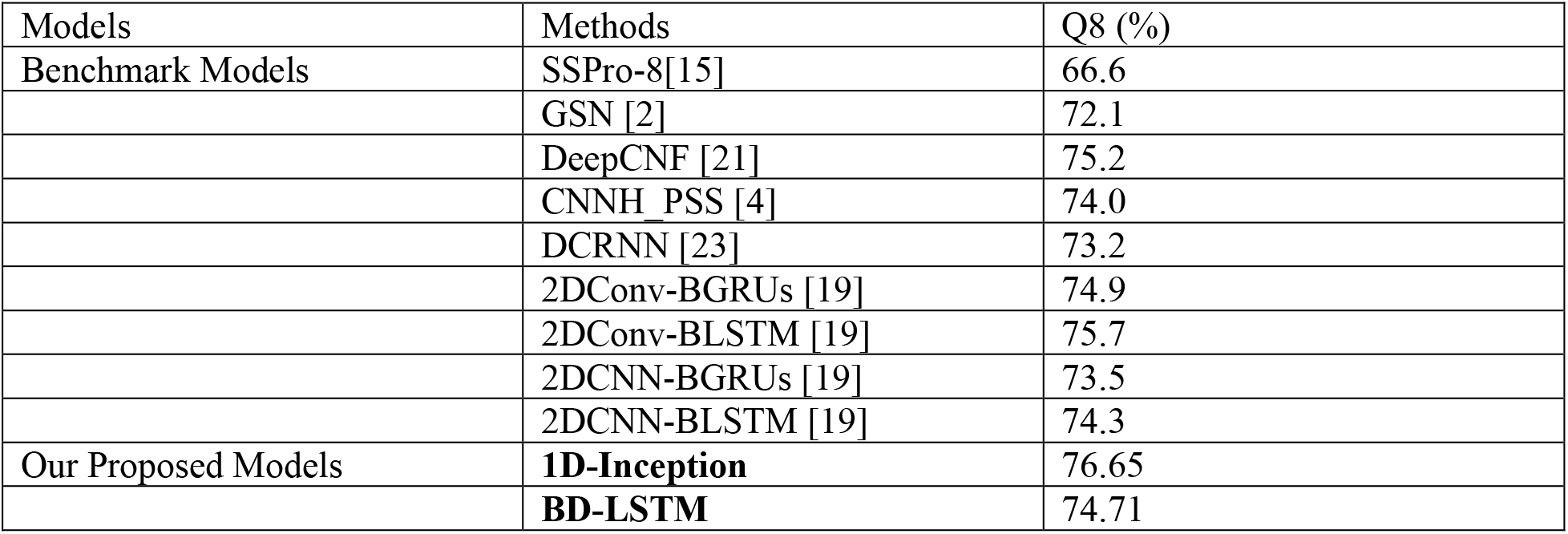
The overall Q8 (%) accuracy for PSS of our proposed models and some existing state-of-art methods on CullPdb 6133 dataset

### Training on CullPDB 6133 filtered dataset and testing on CB513, CASP10, and CASP11 dataset

In this section, we compare several benchmark models with our proposed models by state-8 (Q8) accuracy of PSS. All models trained on CULLPdb6133 filtered and tested on CB513, CASP10, and CASP11 datasets.

According to table 4, our proposed architecture 1D-Inception present 71.18%, 76.86%, 74.07% Q8 accuracy for CB513, CASP10, and CASP11 datasets respectively. 1D-Inception displays superior overall accuracy than any other state-of-the-art models for protein secondary structure on CB513 and CASP10. It almost provides the same accuracy as MUFOLD-SS (only .44% less) [8] on the CASP11 dataset. For CB513, and CASP10 dataset 1D-Inception provides 0.55%, and .39% higher Q8 accuracy than MUFOLD-SS, respectively. Furthermore, we get .68%, 1.86%, and 1.07% respectively higher accuracy than DeepACLSTM on CB513, CASP10, and CASP11. BD-LSTM also provides promising outcomes than most of the existing benchmark models.

**Table 4:** The overall Q8 (%) accuracy for PSS of our proposed models and some benchmark methods on the CB513, CASP10, and CASP11 dataset

| Models | Methods | CB513(%) | CASP10(%) | CASP11(%) |
| --- | --- | --- | --- | --- |
| Benchmark Models | SSPro-8[15] | 63.5 | 64.9 | 65.6 |
|  | GSN [2] | 66.4 | - | - |
|  | CNF [20] | 64.9 | 64.8 | 65.1 |
|  | DeepCNF [21] | 68.3 | 71.8 | 72.3 |
|  | CNNH_PSS [4] | 70.3 | - | - |
|  | DCRNN [22] | 69.4 | - | - |
|  | DeepACLSTM [18] | 70.5 | 75.0 | 73.0 |
|  | MUFOLD-SS [8] | 70.63 | 76.47 | 74.51 |
|  | 2DConv-BGRUs[19] | 69.0 | 72.2 | 71.0 |
|  | 2DConv-BLSTM[19] | 70.2 | 74.5 | 72.5 |
|  | 2DCNN-BGRUs[19] | 68.7 | 72.1 | 71.7 |
|  | 2DCNN-BLSTM[19] | 70.0 | 74.5 | 72.6 |
| Our Proposed Models | **1D-Inception** | 71.18 | 76.86 | 74.07 |
|  | **BD-LSTM** | 69.49 | 74.07 | 72.37 |

In our knowledge, 1D-Inception provides superior Q8 accuracy on CullPdb 6133, CB513, and CASP10 than any other existing method and nearly as excellent Q8 efficiency as MUFOLD-SS on CASP11.

## Conclusion and Discussion

Right now, protein secondary structure prediction is an immensely important task in computational biology because of drug discovery, drug design, and analyzing protein function. In this paper, we proposed two different new deep learning architectures to predict protein secondary structure accurately. To be specific, we exploit the deep convolutional neural network (DCNN) and recurrent neural network (RNN) to find the state-8 PSS. Here, five well-known datasets used for training, validation, and testing these new methods. The deep CNN based approach 1D-Inception exhibits superior performance compared to most of the existing state-of-the-art models on four different datasets: CullPdb6133, CB513, CASP10, and CASP11. To be specific, 1D-Inception outperforms other state-of-the-art methods on CullPdb6133, CB513, and CASP10. Additionally, our RNN based architecture BD-LSTM also produces promising accuracy than many recent approaches to predict PSS.

Compared to the past deep learning models for 8-categories of PSS prediction, 1D-Inception uses a sophisticated approach, but it is efficient to predict secondary structure. Most of the existing architecture cannot extract long-range interdependencies or local context properly. However, 1D-Inception appropriately processes the local and long-distance interdependencies between amino acid residues in the protein sequence, which are one of the significant reasons to enhance the accuracy of the 8-classes protein secondary structure.

Our proposed models can also be utilized in other sequence labeling problems, and it is not limited in computational biology. In the future, we will try to reduce the computational complexities and time complexities of these models so that it can be useful for the practical world. Additionally, adding some physical properties in the input features of proteins to increase the prediction accuracy. Lastly, we will like to add attention mechanisms in our inception modules for the balance-related problem and low-frequency global interactions of protein secondary structure prediction.

